# SARS-CoV-2 variants of concern remain dependent on IFITM2 for efficient replication in human lung cells

**DOI:** 10.1101/2021.11.17.468942

**Authors:** Rayhane Nchioua, Annika Schundner, Dorota Kmiec, Caterina Prelli-Bozzo, Fabian Zech, Lennart Koepke, Alexander Graf, Stefan Krebs, Helmut Blum, Manfred Frick, Konstantin M. J. Sparrer, Frank Kirchhoff

## Abstract

It has recently been shown that an early SARS-CoV-2 isolate (NL-02-2020) hijacks interferon-induced transmembrane proteins (IFITMs) for efficient replication in human cells. To date, several “Variants of Concern” (VOCs) showing increased infectivity and resistance to neutralization have emerged and globally replaced the early viral strains. Here, we determined whether the four SARS-CoV-2 VOCs (Alpha, Beta, Gamma and Delta) maintained the dependency on IFITM proteins for efficient replication. We found that depletion of IFITM2 strongly reduces viral RNA production by all four VOCs in the human epithelial lung cancer cell line Calu-3. Silencing of IFITM1 had little effect, while knock-down of IFITM3 resulted in an intermediate phenotype. Strikingly, depletion of IFITM2 generally reduced infectious virus production by more than four orders of magnitude. In addition, an antibody directed against the N-terminus of IFITM2 inhibited SARS-CoV-2 VOC replication in iPSC-derived alveolar epithelial type II cells thought to represent major viral target cells in the lung. In conclusion, endogenously expressed IFITM proteins (especially IFITM2) are critical cofactors for efficient replication of genuine SARS-CoV-2 VOCs, including the currently dominating Delta variant.

**IMPORTANCE:** Recent results showed that an early SARS-CoV-2 isolate requires endogenously expressed IFITM proteins for efficient infection. However, whether IFITMs are also important cofactors for infection of emerging SARS-CoV-2 VOCs that out-competed the original strains and currently dominate the pandemic remained to be determined. Here, we demonstrate that depletion of endogenous IFITM2 expression almost entirely prevents the production of infectious Alpha, Beta, Gamma and Delta VOC SARS-CoV-2 virions in a human lung cell line. In comparison, silencing of IFITM1 had little impact, while knock-down of IFITM3 had intermediate effects on viral replication. Finally, an antibody targeting the N-terminus of IFITM2 inhibited SARS-CoV-2 VOC replication in iPSC-derived alveolar epithelial type II cells. Our results show that SARS-CoV-2 VOCs including the currently dominant Delta variant are dependent on IFITM2 for efficient replication suggesting that IFITM proteins play a key role in viral transmission and pathogenicity.

## INTRODUCTION

Since its first occurrence in Wuhan (China) in December 2019, the severe acute respiratory syndrome coronavirus 2 (SARS-CoV-2), the causative agent of coronavirus disease 2019 (COVID-19), has caused a devastating pandemic (1, 2). The reasons for the efficient spread of this coronavirus are not fully understood but clearly involve the ability to efficiently infect and propagate in human cells. Viral entry depends on binding of the viral Spike (S) protein to the cellular angiotensin-converting enzyme (ACE) 2 receptor and proteolytic processing of the S precursor into the active S1 and S2 subunits (3–5). However, additional host factors may affect the efficiency of SARS-CoV-2 entry and play roles in viral transmission and pathogenesis (6).

We recently demonstrated that interferon-inducible transmembrane proteins (IFITMs 1, 2, and 3) are required for efficient SARS-CoV-2 infection (7). This came as surprise since IFITMs are a family of IFN stimulated genes (ISGs) that are well-known to protect cells against infection by numerous viral pathogens including retro-, flavi-, influenza-, rhabdo-, filo- and bunyaviruses (3–5). Inhibitory effects have also been reported for highly pathogenic coronaviruses, including SARS-CoV-2 (8,9). However, most evidence was obtained using pseudo-particles containing the S protein of SARS or MERS coronaviruses and/or cells that are not intrinsically permissve to this virus, artificially overexpress IFITM proteins and. Notably, it has been reported that the common cold coronavirus OC43 hijacks IFITM3 for efficient entry (10).

The antiviral mechanism of IFITMs is thought to involve alterations in the rigidity and curvature of the cellular membrane, affecting viruses in a broad, unspecific way (4, 5, 11). In contrast, the SARS-CoV-2 enhancing effect most likely involves specific interactions between the S protein and the N-terminal region of IFITMs, especially IFITM2 (7), suggesting that this pandemic viral pathogen hijacks IFITMs for efficient infection. In accordance with this knockdown of endogenous IFITM2 expression in human lung cell lines strongly reduced viral entry and infectious virus production. In addition, IFITM2-derived peptides as well as an IFITM2-targeting antibody protected gut organoids and cardiomyocytes against infection and cytopathic effects of SARS-CoV-2 (7).

In the initial study, IFITM dependency for efficient infection has only been demonstrated for an early European variant of SARS-CoV-2 isolated in the Netherlands in February 2020 (NL-02-2020) (7). Since then numerous variants have emerged. Some of them show increased transmission fitness and immune evasion and are thus referred to as “variants of concern” (VOCs). Currently, the WHO has categorized five SARS-CoV-2 variants as VOCs: B.1.1.7, B.1.351, P.1, B.1.617.2 and B.1.1.529. The first four, referred to as Alpha, Beta, Gamma and Delta variants, have significantly spread in the human population. The latest (Omicron) variant contains a worrisome number of mutations (12). However, it remains to be seen whether it might outcompete the Delta variant that currently dominates the pandemic and the Omicron variant was not yet Available for biological characterization. Compared to the NL-02-2020 isolate, all VOCs contain various alterations in their S proteins reported to enhance viral infectivity, transmissibility and pathogenicity by affecting ACE2 receptor affinity, proteolytic activation and susceptibility to neutralization (13–15). Here, we examined whether current SARS-CoV-2 VOCs still require IFITM proteins for efficient replication in human lung cells.

## RESULTS

To verify that the SARS-CoV-2 VOCs show the expected differences in S compared to the early NL-02-2020 isolate, we performed full-genome sequence analyses of the four variants used for functional analyses. The NL-02-2020 isolate already contains the D614G amino acid substitution within the receptor-binding motif (RBM) which has been reported to increase SARS-CoV-2 transmissibility by promoting ACE2 receptor interaction and is also found in all current VOCs (16, 17). As expected, the Spike proteins of the VOCs differed by six to ten amino acid changes from the NL-02-2020 Spike (Figure 1). The Alpha VOC (B.1.1.7) that emerged at the end of 2020 in the UK contains eight mutations in its S protein: Deletion of H69/V70 and Y144; Mutation of N501Y, A570D, P681H, T716I, S982A and D1118H (18). The Beta VOC (B.1.351) emerged in South Africa in October 2020 and has initially spread to all continents (19). Its S protein contains three alterations in the RBD (K417N, E484K, N501Y) and five additional changes (L18F, D80A, D215G, R246I, A701V). The Gamma (P.1) variant was first detected in Brazil at the end of 2020 and shares the K417T, E484K and N501Y S mutations with the Alpha and/or Beta VOCs (Figure 1) (20). The Delta (B.1.617.2) variant was first identified in India in the first half of 2021 (21) and has efficiently outcompeted all other SARS-CoV-2 VOCs around the globe. It differs by changes of T19R, deletion of residues 157-158, L452R, T478K, E484K, P681R, R683L and D950N from NL-02-2020 in the S protein (Figure 1). Several changes (L18F, K417T, E484K and N501Y) emerged independently by convergent evolution in several VOCs (14, 22). The reasons why they are associated with a selective advantage remain to be fully elucidated but rapidly accumulating evidence shows they reduce neutralization by antibodies and/or increase ACE2 binding affinity (14). In addition, the P681R substitution near the furin cleavage site might improve proteolytic activation of the Delta S protein (23, 24). Thus, all VOCs contain changes in their S proteins reported to increase interaction with their primary ACE2 receptor that might also affect the dependency on other cellular cofactors for efficient entry and fusion, like IFITM proteins.

**Figure 1:**
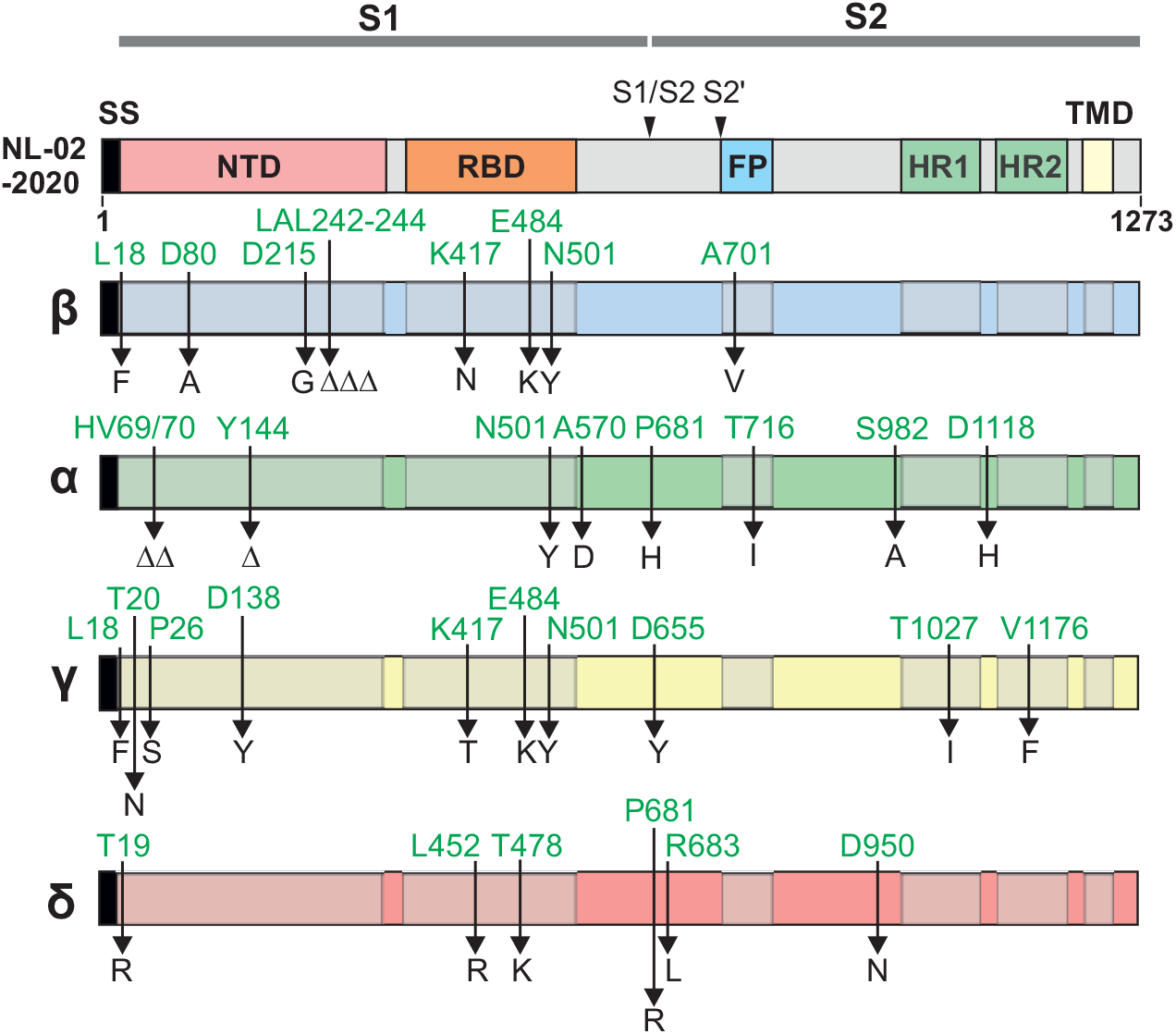
Amino acid variations in the Spike proteins of SARS-CoV-2 variants investigated. The upper panel shows a schematic representation of the SARS-CoV-2 S protein with specific domains indicated in different colours. Abbreviations: SS, signal sequence; NTD, N-terminal domain; RBD, receptor binding domain; FP, fusion peptide; HR, heptad repeat and TMD, transmembrane domain. The S1/S2 and S2’ proteolytic cleavage sites are also indicated.

To examine the role of endogenous IFITM expression on infection by genuine SARS-CoV-2 VOCs, we performed siRNA knockdown (KD) studies in the human epithelial lung cancer cell line Calu-3, which endogenously express ACE2 and all three IFITM proteins (7). Viral replication was determined by quantification of viral N (nucleocapsid) RNA levels by qRT-PCR in the cell culture supernatants 2 days after infection with the five SARS-CoV-2 variants (Figure 2A). Most VOCs produced 2-to 4-fold higher levels of viral RNA compared to NL-02-2020 in Calu-3 cells (Figure 2B). The single exception was the Beta variant, which showed moderately lower levels of viral RNA production. Silencing of IFITM2 expression reduced viral RNA production from 31-(Alpha) to 754-fold (Gamma). Replication of the Beta variant was 112x reduced in the absence of IFITM2, respectively. In comparison, KD of IFITM1 had little effect, while silencing of IFITM3 resulted in 2-(Beta) to a maximum of 31-fold (NL-02-2020) lower viral RNA yields (Figure 2B). Notably, IFITM2 KD still reduced viral RNA yields by the currently dominant Delta variant by >100-fold, while IFITM3 silencing was associated with a 20-fold reduction, demonstrating that this VOC still requires IFITM proteins for efficient entry.

**Figure 2:**
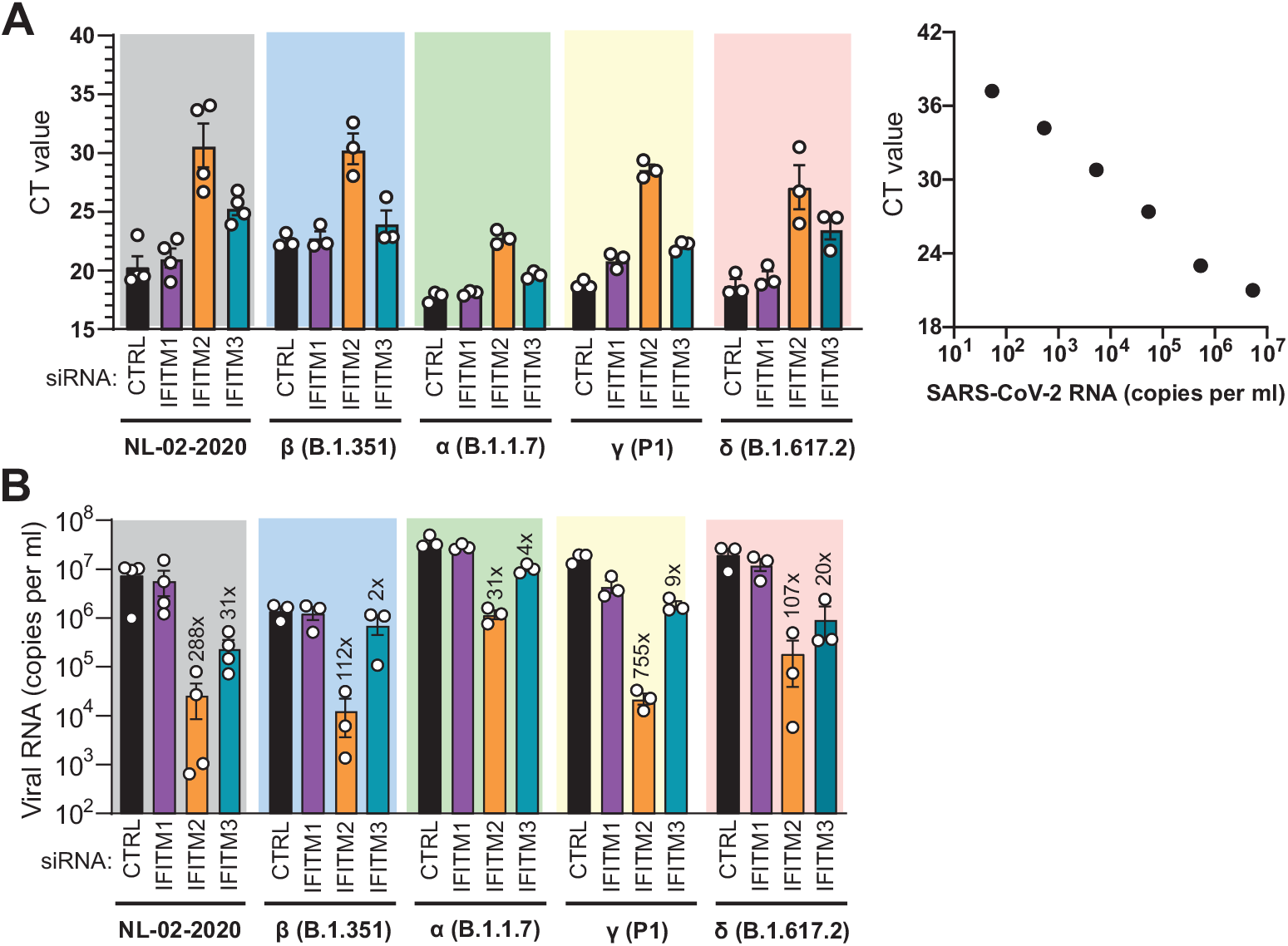
Role of IFITMs in replication of SARS-CoV-2 VOCs in Calu-3 cells. (**A**) Standard curve and raw qRT-PCR CT values obtained using supernatants of Calu-3 cells collected 2 days post-infection. (**B**) Viral N RNA levels in the supernatant of Calu-3 cells infected with the indicates SARS-CoV-2 variants. Cells were transfected with control (CTRL) or IFITM targeting siRNAs as indicated. Numbers above the bars indicate n-fold reduction compared to the viral RNA levels detected in the supernatant of Calu-3 cells treated with CTRL siRNA. Bars in panel A and B represent the mean of 3 to 4 independent experiments (±SEM) each measured in technical duplicates.

To further determine whether IFITM2 is critical for productive replication of SARS-CoV-2 VOCs in Calu-3 cells, we determined the TCID_50_ (50% Tissue Culture Infectious Dose) of viral particles in the cell culture supernatants (Figure 3A). With the exception of the Beta variant that showed the lowest viral RNA yields (Figure 2) and infectious titers, all SARS-CoV-2 variants produced more than 10 million infectious virus particles per ml culture supernatant in Calu-3 cells treated with the control siRNA (Figure 3B). In striking contract, infectious virus production was generally reduced to levels near or below background (≤100 infectious particles per ml) upon silencing of IFITM2 (Figure 3). Altogether, these results show that all four SARS-CoV-2 VOCs including the dominant Delta variants are strongly dependent on endogenous IFITM2 expression for efficient replication in Calu-3 cells.

**Figure 3:**
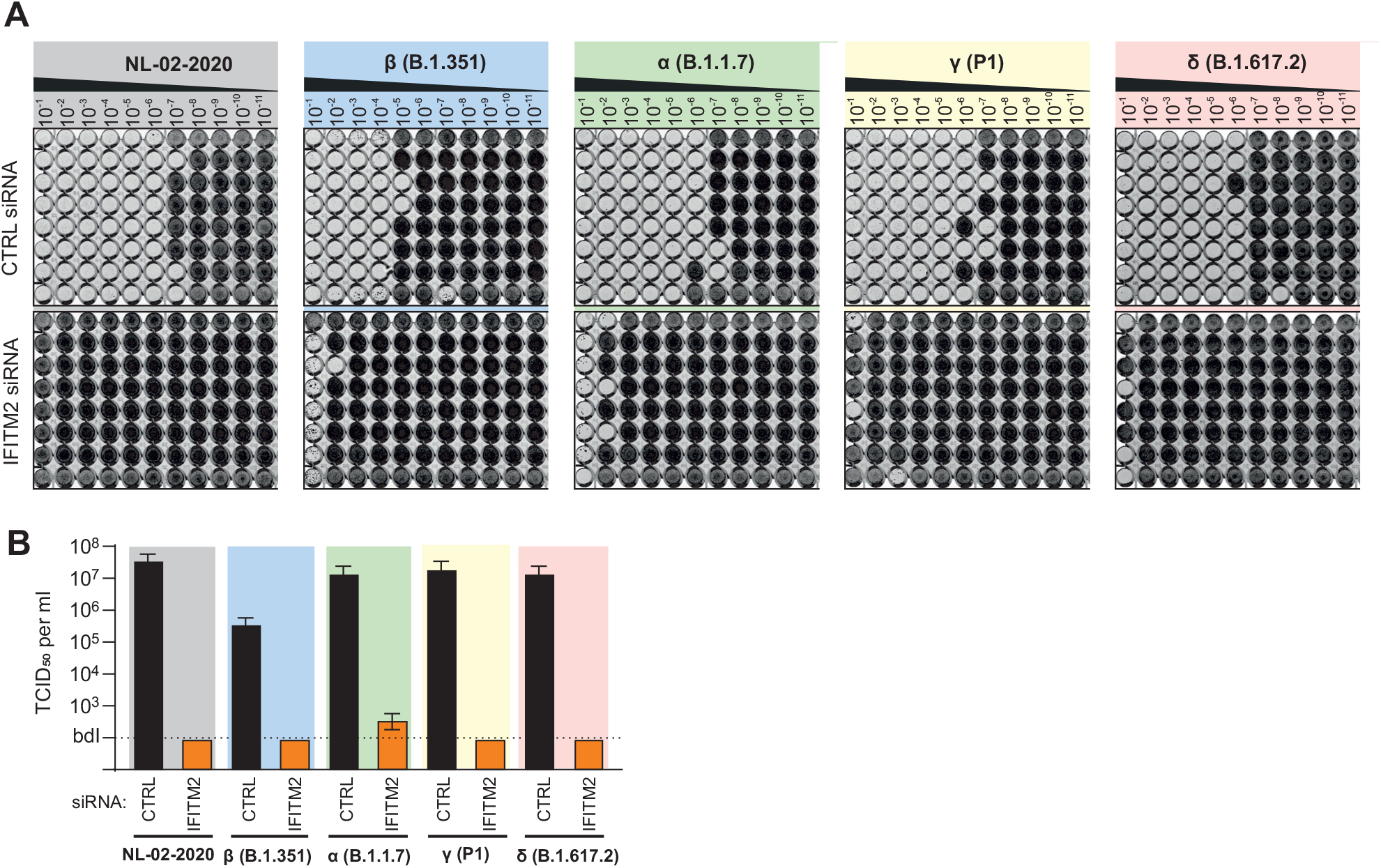
Silencing of endogenous IFITM2 expression prevents production of infectious SARS-CoV-2. (**A**) Supernatants derived from Calu-3 cells treated with control (CTRL) or IFITM2 siRNA two days after infection with the SARS-CoV-2 NL-02-2020 or the indicated VOCs were serially diluted and added to Caco-2 cells seeded in 96-well plates. Five days later, cells were examined for CPE, fixed and stained with crystal violet. Productively infected wells appear transparent since the cells are eliminated or detached due to viral infection. (**B**) Quantification of infectious SARS-CoV-2 particles in the supernatant of Calu-3 cells treated with control or IFITM2 targeting siRNAs. Bars represent the mean of one experiment performed with eight technical replicates (±SD) shown in panel A. Bdl, below detection limit.

We have previously shown that an antibody targeting the N-terminus of IFITM2 inhibits replication of the NL-02-2020 isolate in gut organoids and cardiomyocytes (7). To further examine the potential relevance of IFITM2 for transmission of SARS-CoV-2 VOCs, we performed experiments in iPSC-derived alveolar epithelial type II (iATII) cells, as a model for the main target cells of SARS-CoV-2 infection in the distal lung (25). Western blot analyses showed that, similarly to Calu-3 cells, iATII cells express IFITM2 and IFITM3 (Figure 4A). In contrast, both cells types showed little (Calu-3) or no (iATII) detectable expression of IFITM1. Unexpectedly, we did not detect ACE2 expression in iATII cells by western blot analyses while ACE2 was readily detectable in Calu-3 cells (Figure 4A). In agreement with published data (26), however, ACE2 expression by iATII cells was clearly detectable by FACS (Figure 4B).

**Figure 4:**
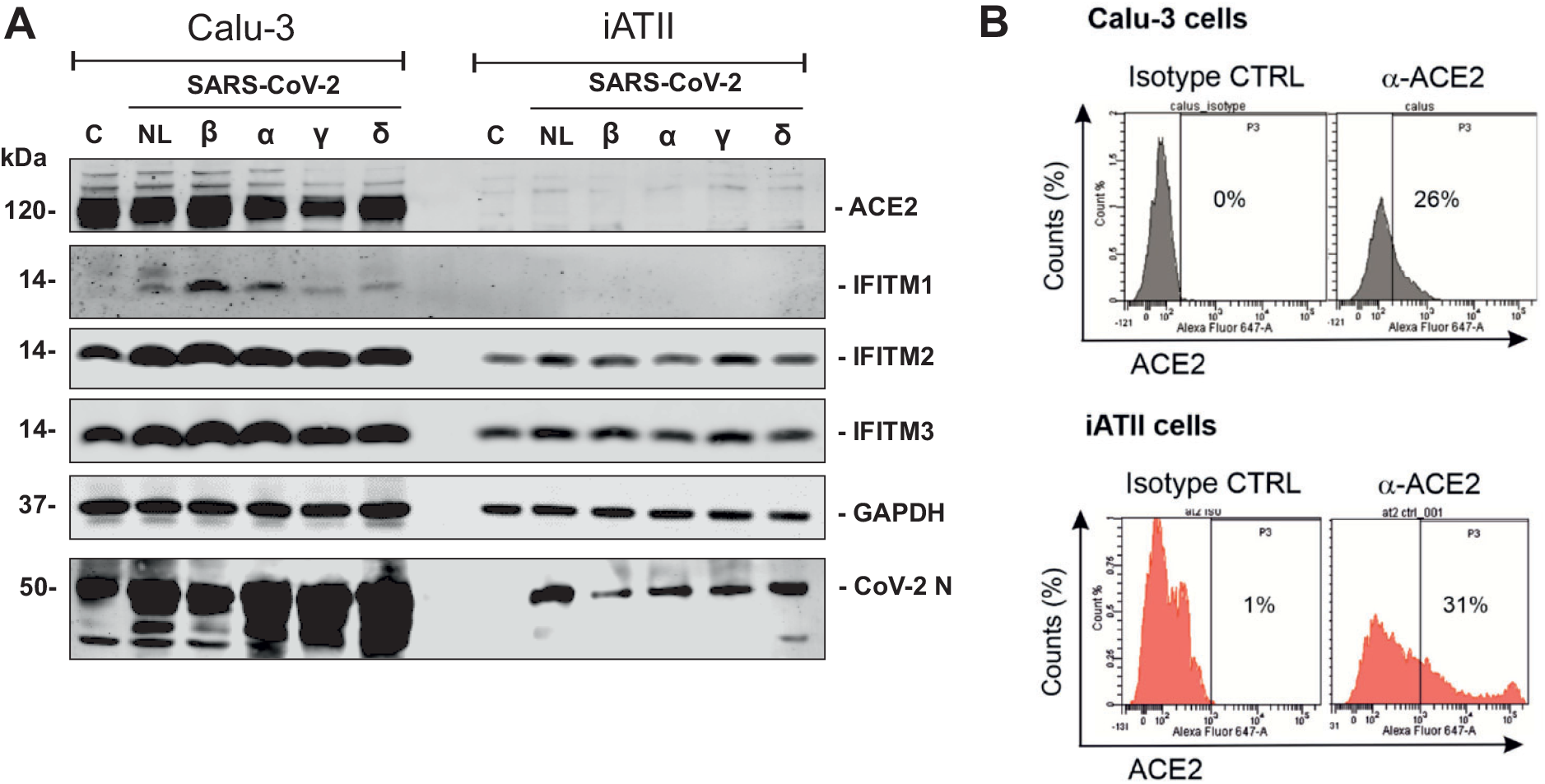
Expression of ACE2 and IFITM proteins in Calu-3 and iATII cells. (**A**) Immunoblot of ACE2, IFITM1, IFITM2, and IFITM3 in Calu-3 and iATII cells left uninfected (c) or infected with the indicated SARS-CoV-2 variants. Whole-cell lysates were stained with the indicated antibodies. An unspecific signal was observed in the Calu-3 control lane stained with the CoV-2 N antibody. (**B**) Flow cytometric analysis of surface ACE2 expression in Calu-3 and iATII cells.

In agreement with published data (26–28), iATII cells were highly susceptible to SARS-CoV-2 replication and typically produced about 100-fold higher levels of viral RNA compared to Calu-3 cells (Figures 2B, 5A). On average, the Delta variant replicated to about 30-fold higher levels (average vRNA copy numbers of 2.4×10^11^) than the early NL-02-2020 isolate in iATII cells (Figure 5A). The broad-spectrum antiviral agent Remdesivir (29) efficiently inhibited replication of all SARS-CoV-2 variants. Treatment of iATII cells with the antibody against the N-terminus of IFITM2 also generally reduced viral RNA production in a dose-dependent manner, albeit varying efficiencies. Most notably, the anti-IFITM2 Ab reduced replication of the Delta VOC in iATII cells by up to 95% (Figure 5B). This agrees with our previous finding that IFITM2 can be targeted to protect various types of human cells against SARS-CoV-2 infection (7).

**Figure 5:**
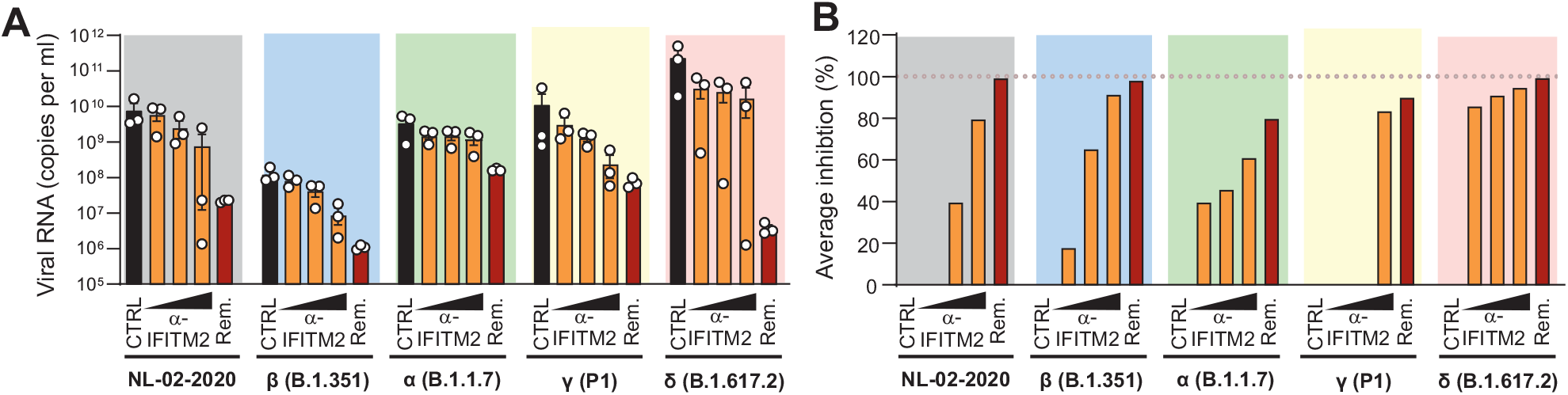
Effect of an *α*-IFITM2 antibody on replication of SARS-CoV-2 variants in iATII cells. (**A**) Quantification of viral N RNA levels in the supernatant of iATII cells treated with α-IFITM2 antibody (20, 40, or 80 µg/ml) or Remdesivir (10 µM) 1 h before infection (SARS-CoV-2, MOI 0.5), collected 48 h post-infection. Bars represent the mean of three independent experiments. (**B**) Average percentage of reduction of vRNA levels in the supernatants of (E) compared to the untreated control.

## DISCUSSION

In the present study, we demonstrate that IFITMs (especially IFITM2) are also critical cofactors for efficient replication of all four SARS-CoV-2 VOCs including the currently dominant Delta variant. We have previously shown that IFITMs also promote SARS-CoV-2 replication in primary small airway epithelial cells (SAEC) cells and that IFITM2 can be targeted to inhibit viral replication in gut organoids and cardiomyocytes derived from human iPSCs (7). The present finding that an *α*-IFITM2 antibody inhibits replication of the Delta variants in iPSC-derived alveolar epithelial type II cells, proposed to model main target cells of SARS-CoV-2 infection in the distal lung (26, 28), adds to the evidence that IFITM2 may play a key role in SARS-CoV-2 transmission, dissemination and pathogenesis. Our observation that IFITM2 dependency is maintained by VOCs also further underlines that against the odds this cellular “antiviral” factor represents a target for therapeutic or preventive approaches.

In agreement with previous findings (23, 30), the Delta variant replicated with higher efficiency than the early SARS-CoV-2 isolate in human lung cells particularly in iPSC-derived alveolar epithelial type II cells. This agrees with recent data showing that the Delta variant infects human bronchial epithelial cells with higher efficiency than other VOCs (31). Altogether, our results show that IFITMs (especially IFITM2) are also critical cofactors for efficient replication of all four current SARS-CoV-2 VOCs. Notably, however, the Alpha variant yielded ∼100-fold higher levels of vRNA upon silencing of IFITM2 expression compared to the 2019 CoV-2 and Beta variants (Figure 2). Thus, it is tempting to speculate that this VOC did not only evolve reduced susceptibility to IFN inhibition (30, 32) but may also dependent less on IFITM2 for efficient infection compared to other SARS-CoV-2 variants.

The exact mechanism of IFITM2-dependent enhancement needs further studies. However, our previous studies suggest that IFITM2 promotes SARS-CoV-2 entry by direct interaction with the viral Spike protein and enhancing virus-cell fusion in early endosomes (7). The enhancing effect is only observed for genuine SARS-CoV-2 and endogenous IFITMs, while the broad antiviral activity of IFITMs involving alterations in cellular membrane rigidity and curvature instead of specific interactions with viral glycoproteins has been reported for pseudovirions carrying SARS-CoV-2 Spike proteins and for overexpression of IFITM proteins (8). In agreement with our previous results (7), IFITM2 KD had stronger effects on infectious titers than on viral RNA yields. It remains to be determined whether the presence of IFITM2 has indeed an enhancing effect on the infectiousness of SARS-CoV-2 particles or if the background levels are just higher for viral RNA due to release from or lysis of infected cells.

The Alpha and Delta variants contain a mutation P681H/R close to the furin cleavage site that might affect interferon sensitivity, proteolytic activation and fusogenicity of the S protein (33, 34). These two VOCs produced the highest levels of viral RNA upon IFITM2 KD and it will be interesting to further examine whether an increased intrinsic fusogenic activity of the Alpha and Delta Spike proteins affects their dependency on IFITM2 for infection.

Our results are further evidence that IFITM proteins are critical cofactors for infection of SARS-CoV-2 in primary human target cells. We show that an *α*-IFITM2 antibody inhibits replication of the Delta VOC in human alveolar epithelial type II cells by >90% (Figure 5B) reported to play a key role in the spread of SARS-CoV-2. This finding further suggests that IFITM2 may be a highly unexpected suitable target for therapeutic approaches against this pandemic viral pathogen. It will be interesting to determine whether the emerging Omicron variant that contains a striking number of about 30 amino acid changes in the Spike protein compared to the Wuhan strain (15, 35) is also dependent on IFITM2 for efficient infection and susceptible to inhibition by *α*-ITITM2 antibodies.

## ACKNOWLEDGMENTS

We thank Daniela Krnavek, Martha Mayer, Kerstin Regensburger, Regina Burger, Jana Romana Fischer and Birgit Ott for laboratory assistance. We also thank Stefan Krebs, Helmut Blum, Alexander Graf for next generation sequencing of SARS-CoV-2 VOCs, Michael Schindler for providing the B.1.351 (Beta) VOC and Florian Schmidt, Beate M. Kümmerer and Hendrick Streeck for providing the B.1.617.2 (Delta) VOC. The following reagent was obtained through BEI Resources, NIAID, NIH: SARS-CoV-2, Isolate hCoV-19/Japan/TY7-503/2021 (Brazil P.1), NR-54982, contributed by National Institute of Infectious Diseases.

## AUTHORS INFORMATION

Conceptualization and funding acquisition, F.K., K.M.J.S., M.F.; Investigation, R.N., C.P.B., F.Z., L.K., D.K. and A. S.; Writing, F.K.; Review and editing, all authors.

## MATERIAL AND METHODS

### Cell culture

Calu-3 (human epithelial lung adenocarcinoma) cells were cultured in Minimum Essential Medium Eagle (MEM, Sigma, Cat#M4655) supplemented with 10% (upon and after viral infection) or 20% (during all other times) heat-inactivated fetal bovine serum (FBS, Gibco, Cat#10270106), 100 units/ml penicillin, 100 µg/ml streptomycin (ThermoFisher, Cat#15140122), 1 mM sodium pyruvate (Pan Biotech, Cat#P04-8010), and 1x non-essential amino acids (Sigma, Cat#M7145). Vero E6 cells (Cercopithecus aethiops derived epithelial kidney, ATCC) and TMPRSS2-expressing Vero E6 cells (kindly provided by the National Institute for Biological Standards and Control (NIBSC), No. 100978) were grown in Dulbecco’s modified Eagle’s medium (DMEM, Gibco, Cat#41965039) supplemented with 2.5% (upon and after viral infection) or 10% (during all other times) heat-inactivated FBS (Gibco, Cat#10270106), 100 units/ml penicillin, 100 µg/ml streptomycin (ThermoFisher, Cat#15140122), 2 mM L-glutamine (Gibco, Cat#25030081), 1 mM sodium pyruvate (Pan Biotech, Cat# P04-8010), 1x non-essential amino acids (Sigma, Cat#M7145) and 1 mg/mL Geneticin (Gibco, Cat#10131-019) (for TMPRSS2-expressing Vero E6 cells). Caco-2 cells (human epithelial colorectal adenocarcinoma, kindly provided by Prof. Holger Barth (Ulm University)) were grown in the same media as Vero E6 cells but with supplementation of 10% heat-inactivated FBS.

Human induced Alveolar Type 2 cells (iATII) were differentiated from BU3 NKX2-1^GFP^;SFTPC^tdTomato^ induced pluripotent stem cells(36) (iPCSs, kindly provided by Darrell Kotton, Boston University and Boston Medical Center) and maintained as alveolospheres embedded in 3D Matrigel in CK+DCI media, as previously described(37). For infection studies, iATII cells were cultured as 2D cultures on Matrigel-coated plates in CK+DCI medium + 10 µM Y-27632 (Tocris, Cat#1254) for 48h before switching to CK+DCI medium on day 3.

### SARS-CoV-2 stocks

The SARS-CoV-2 variant B.1.351 (Beta), 2102-cov-IM-r1-164 was provided by Prof. Michael Schindler (University of Tübingen) and the B.1.617.2 (Delta) variant by Prof. Florian Schmidt (University of Bonn). The BetaCoV/Netherlands/01/NL/2020 (NL-02-2020) and B.1.1.7. (Alpha) variants were obtained from the European Virus Archive. The hCoV-19/Japan/TY7-503/2021 (Brazil P.1) (Gamma) (#NR-54982) isolate was obtained from the BEI resources. SARS-CoV-2 strains were propagated on Vero E6 (NL-02-2020, Delta), VeroE6 overexpressing TMPRSS2 (Alpha), CaCo-2 (Beta) or Calu-3 (Gamma) cells. To this end, 70-90% confluent cells in 75 cm^2^ cell culture flasks were inoculated with the SARS-CoV-2 isolate (multiplicity of infection (MOI) of 0.03-0.1) in 3.5 ml serum-free medium. The cells were incubated for 2h at 37°C, before adding 20 ml medium containing 15 mM HEPES (Carl Roth, Cat#6763.1). Virus stocks were harvested as soon as strong cytopathic effect (CPE) became apparent. The virus stocks were centrifuged for 5 min at 1,000 g to remove cellular debris, aliquoted, and stored at -80°C until further use.

### Sequencing of SARS-CoV-2 VOCs

Virus stocks were inactivated and lysed by adding 0.3 ml TRIzol Reagent (ambion, Cat#132903) to 0.1 ml virus stock. Viral RNA was isolated using the Direct-zol RNA MiniPrep kit (ZymoResearch, Cat#R2050) according to manufacturer’s instructions, eluting the RNA in 50 µl DNase/RNase free water. The protocol to prepare the viral RNA for sequencing was modified from the nCoV-2019 sequencing protocol V.1. For reverse transcription, the SuperScript IV First-Strand Synthesis System (Invitrogen, Cat#18091050) was used with modified manufacturer’s instructions. First, 1 µl random hexamers (50 ng/µl), 1 µl dNTPs mix (10 mM each), and 11 µl template RNA (diluted 1:10 in DNase/RNase free water) were mixed, incubated at 65°C for 5 min and placed on ice for 1 min. Next, 4 µl SSIV Buffer, 1 µl DTT (100 mM), 1 µl RNaseOUT RNase Inhibitor, and 1 µl SSIV Reverse Transcriptase were added to the mix, followed by incubation at 24°C for 5 min, 42°C for 50 min, and 70°C for 10 min. To generate 400 nt fragments in PCR, the ARTIC nCoV-2019 V3 Primer set (IDT) and the Q5 Hot Start High-Fidelity 2X Master Mix (NEB, Cat#M0494S) were used with modified manufacturer’s instructions. The primers pools 1 and 2 were diluted to a final concentration of 10 µM and a reaction with each primer pool was set up as follows, 4 µl respective primer pool, 12.5 µl Q5 Hot Start High-Fidelity 2X Master Mix, 6 µl water, and 2.5 µl cDNA. The PCR was performed as follows, 98°C for 30 s, 30 cycles of 98°C for 15 s and 65°C for 5 min, and hold at 4°C. The PCR products were run on a 1% agarose gel to check for the presence of fragments at the correct size. The products from primer pool 1 and primer pool 2 for each variant were pooled, diluted and quantified by Qubit DNA HS kit (Thermo Fisher, Cat#Q32851). The sequencing amplicon pools were diluted to 0.2 ng/µl and tagmented with Nextera XT library prep kit (Illumina, Cat#FC-131-1024). Nextera libraries were dual-barcoded and sequenced on an Illumina NextSeq1000 instrument. The obtained sequenced reads were demultiplexed and mapped against the SARS-CoV-2 reference genome (NC_045512.2) with *BWA-MEM*(38). Pileup files were generated from the mapped reads using *Samtools*(39). The mapped reads and the pileup file were used to construct the consensus sequence with the *iVar* package (40) using default settings.

### IFITM knock-down

At 24 h and 96 h post-seeding, 150,000 Calu-3 cells, seeded in 24-well plates, were transfected with 20 µM of non-targeting siRNA or IFITM1, IFITM2 or IFITM3 specific siRNAs using Lipofectamine RNAiMAX (Thermo Fisher, Cat#13778100) according to the manufacturer’s instructions. 6 h after the second transfection, Calu-3 cells were infected with the various SARS-CoV-2 variants at an MOI of 0.05. 6 h post-infection, the inoculum was removed, the cells washed once with PBS and supplemented with fresh media. 48 h post infection, supernatants were harvested for qRT-PCR analysis.

### qRT-PCR

*N* (nucleoprotein) transcript levels were determined in supernatants collected from SARS-CoV-2 infected Calu-3 cells 48 h post-infection as previously described(41). Total RNA was isolated using the Viral RNA Mini Kit (Qiagen, Cat#52906) according to the manufacturer’s instructions. RT-qPCR was performed as previously described(42) using TaqMan Fast Virus 1-Step Master Mix (Thermo Fisher, Cat#4444436) and a OneStepPlus Real-Time PCR System (96-well format, fast mode). Primers were purchased from Biomers (Ulm, Germany) and dissolved in RNAse free water. Synthetic SARS-CoV-2-RNA (Twist Bioscience, Cat#102024) or RNA isolated from BetaCoV/France/IDF0372/2020 viral stocks quantified via this synthetic RNA (for low Ct samples) were used as a quantitative standard to obtain viral copy numbers. All reactions were run in duplicates. Forward primer (HKU-NF): 5’-TAA TCA GAC AAG GAA CTG ATT A-3’; Reverse primer (HKU-NR): 5’-CGA AGG TGT GAC TTC CAT G-3’; Probe (HKU-NP): 5’-FAM-GCA AAT TGT GCA ATT TGC GG-TAMRA-3’.

### Inhibition by IFITM2 antibody and Remdesivir

30,000 iATII cells were seeded as single cells in 96-well plates coated for 1 h at 37 °C with 0.16 mg/ml Matrigel (Corning, Cat#356238) diluted in DMEM/F12 (Thermo Fisher, Cat#11330032), 24 h later cells were treated with increasing concentrations (20, 40, 80 µg/ml) of α-IFITM2 (Cell Signaling, Cat#13530 S) or Remdesivir (Selleck Chemicals Cat#S8932) (10 µM). One h post-treatment, cells were infected with SARS-CoV-2 VOCs with a MOI of 0.5. 6 h post-infection, cells were washed once with PBS and supplemented with fresh medium. Thereafter, day 0 wash CTRL was harvested. 48 h post-infection supernatants were harvested for qRT-PCR analysis.

### Western blot

To determine the expression of cellular and viral proteins, infected Calu-3 (MOI 0.2, 48 h post infection) or iATII (MOI 0.5, 48 h post infection) cells or uninfected controls were washed in PBS and subsequently lysed in Western blot lysis buffer (150 mM NaCl, 50 mM HEPES, 5 mM EDTA, 0.1% NP40, 500 μM Na3VO4, 500 μM NaF, pH 7.5) supplemented with protease inhibitor cocktail (Roche, Cat#11697498001). After 5 min of incubation on ice, samples were centrifuged (4 °C, 20 min, 20,817 g) to remove cell debris. The supernatant was transferred to a fresh tube, the protein concentration was measured by Nanodrop and adjusted using Western blot lysis buffer. Western blotting was performed as previously reported. Proteins were stained using primary antibodies against IFITM1 (α-IFITM1, Cell Signaling Cat#13126S, 1:1,000), IFITM2 (α-IFITM2 Cell Signaling Cat#13530 S, 1:1,000), IFITM3 (α-IFITM3 Cell Signaling Cat#59212S, 1:1,000), ACE2 (Rabbit polyclonal anti-ACE2 Abcam, Cat#ab166755, 1:1,000); rat anti-GAPDH (Biolegend Cat#607902, 1:1,000) and SARS CoV-2 N (anti-SARS-CoV-2 N Sino Biologicals Cat#40588-V08B, 1:1,000) and Infrared Dye labeled secondary antibodies (LI-COR IRDye). Membranes were scanned using an Odyssey infrared imager and band intensities were quantified in Image Studio Lite Version 5.0.

### TCID_50_ Endpoint titration

10,000 Caco-2 cells were seeded in 96-well F-bottom plates. One day later, infectious supernatants were serially diluted and added to the cells. Cells were then incubated for 5 days and monitored for CPE. TCID_50_/mL was calculated according to Reed and Muench.

### Flow cytometric analysis

60,000 iATII cells or Calu-3 cells were incubated for 1 h at 4°C with equal protein concentrations of control rabbit IgG (Diagenode, Cat#C15410206) or 1/200 dilution of rabbit anti-ACE2 (Abcam, Cat#ab166755) diluted in FACS buffer (PBS, 1% FBS), washed three times in PBS, stained for 30 min with 1/400 dilution of goat anti-rabbit AF647 (Invitrogen, Cat#A27040), fixed in 1% PFA and analyzed using a BD FACS Canto II flow cytometer.

### Statistical analysis

Statistical analysis was performed using GraphPad Prism software. Two-tailed unpaired Student’s t-test were used to determine statistical significance. Significant differences are indicated as: *, p < 0.05; **, p < 0.01; ***, p < 0.001; **** p<0.0001. Statistical parameters are specified in the figure legends.

## REFERENCES

1. Zhou P, Yang XL, Wang XG, Hu B, Zhang L, Zhang W, Si HR, Zhu Y, Li B, Huang CL, Chen HD, Chen J, Luo Y, Guo H, Jiang RD, Liu MQ, Chen Y, Shen XR, Wang X, Zheng XS, Zhao K, Chen QJ, Deng F, Liu LL, Yan B, Zhan FX, Wang YY, Xiao GF, Shi ZL. 2020. A pneumonia outbreak associated with a new coronavirus of probable bat origin. Nature 579:270–273.

2. Zhu N, Zhang D, Wang W, Li X, Yang B, Song J, Zhao X, Huang B, Shi W, Lu R, Niu P, Zhan F, Ma X, Wang D, Xu W, Wu G, Gao GF, Tan W, China Novel Coronavirus Investigating and Research Team. 2020. A Novel Coronavirus from Patients with Pneumonia in China, 2019. N Engl J Med 382:727–733.

3. Bailey CC, Zhong G, Huang I-C, Farzan M. 2014. IFITM-Family Proteins: The Cell’s First Line of Antiviral Defense. Annual Review of Virology 1:261–283.

4. Diamond MS, Farzan M. 2013. The broad-spectrum antiviral functions of IFIT and IFITM proteinsNature Reviews Immunology. Nature Publishing Group.

5. Zhao X, Li J, Winkler CA, An P, Guo JT. 2019. IFITM genes, variants, and their roles in the control and pathogenesis of viral infectionsFrontiers in Microbiology. Frontiers Media S.A.

6. Baggen J, Vanstreels E, Jansen S, Daelemans D. 2021. Cellular host factors for SARS-CoV-2 infection. Nat Microbiol 6:1219–1232.

7. Prelli Bozzo C, Nchioua R, Volcic M, Koepke L, Krüger J, Schütz D, Heller S, Stürzel CM, Kmiec D, Conzelmann C, Müller J, Zech F, Braun E, Groß R, Wettstein L, Weil T, Weiß J, Diofano F, Rodríguez Alfonso AA, Wiese S, Sauter D, Münch J, Goffinet C, Catanese A, Schön M, Boeckers TM, Stenger S, Sato K, Just S, Kleger A, Sparrer KMJ, Kirchhoff F. 2021. IFITM proteins promote SARS-CoV-2 infection and are targets for virus inhibition in vitro. Nat Commun 12:4584.

8. Shi G, Kenney AD, Kudryashova E, Zani A, Zhang L, Lai KK, Hall-Stoodley L, Robinson RT, Kudryashov DS, Compton AA, Yount JS. 2020. Opposing activities of IFITM proteins in SARS-CoV-2 infection. EMBO J e106501.

9. Wrensch F, Winkler M, Pöhlmann S. 2014. IFITM proteins inhibit entry driven by the MERS-Coronavirus Spike protein: Evidence for Cholesterol-Independent Mechanisms. Viruses 6:3683–3698.

10. Zhao X, Guo F, Liu F, Cuconati A, Chang J, Block TM, Guo JT. 2014. Interferon induction of IFITM proteins promotes infection by human coronavirus OC43. Proceedings of the National Academy of Sciences of the United States of America 111:6756–6761.

11. Compton AA, Bruel T, Porrot F, Mallet A, Sachse M, Euvrard M, Liang C, Casartelli N, Schwartz O. 2014. IFITM proteins incorporated into HIV-1 virions impair viral fusion and spread. Cell Host and Microbe 16:736–747.

12. Classification of Omicron (B.1.1.529): SARS-CoV-2 Variant of Concern.

13. Soh SM, Kim Y, Kim C, Jang US, Lee H-R. 2021. The rapid adaptation of SARS-CoV-2-rise of the variants: transmission and resistance. J Microbiol 59:807–818.

14. Harvey WT, Carabelli AM, Jackson B, Gupta RK, Thomson EC, Harrison EM, Ludden C, Reeve R, Rambaut A, COVID-19 Genomics UK (COG-UK) Consortium, Peacock SJ, Robertson DL. 2021. SARS-CoV-2 variants, spike mutations and immune escape. Nat Rev Microbiol 19:409–424.

15. Genovese L, Zaccaria M, Farzan M, Johnson WE, Momeni B. 2021. Investigating the mutational landscape of the SARS-CoV-2 Omicron variant via ab initio quantum mechanical modeling.

16. Korber B, Fischer WM, Gnanakaran S, Yoon H, Theiler J, Abfalterer W, Hengartner N, Giorgi EE, Bhattacharya T, Foley B, Hastie KM, Parker MD, Partridge DG, Evans CM, Freeman TM, de Silva TI, Sheffield COVID-19 Genomics Group, McDanal C, Perez LG, Tang H, Moon-Walker A, Whelan SP, LaBranche CC, Saphire EO, Montefiori DC. 2020. Tracking Changes in SARS-CoV-2 Spike: Evidence that D614G Increases Infectivity of the COVID-19 Virus. Cell 182:812–827.e19.

17. Yurkovetskiy L, Wang X, Pascal KE, Tomkins-Tinch C, Nyalile TP, Wang Y, Baum A, Diehl WE, Dauphin A, Carbone C, Veinotte K, Egri SB, Schaffner SF, Lemieux JE, Munro JB, Rafique A, Barve A, Sabeti PC, Kyratsous CA, Dudkina NV, Shen K, Luban J. 2020. Structural and Functional Analysis of the D614G SARS-CoV-2 Spike Protein Variant. Cell 183:739–751.e8.

18. Borges V, Sousa C, Menezes L, Gonçalves AM, Picão M, Almeida JP, Vieita M, Santos R, Silva AR, Costa M, Carneiro L, Casaca P, Pinto-Leite P, Peralta-Santos A, Isidro J, Duarte S, Vieira L, Guiomar R, Silva S, Nunes B, Gomes JP. 2021. Tracking SARS-CoV-2 lineage B.1.1.7 dissemination: insights from nationwide spike gene target failure (SGTF) and spike gene late detection (SGTL) data, Portugal, week 49 2020 to week 3 2021. Euro Surveill 26.

19. Tegally H, Wilkinson E, Giovanetti M, Iranzadeh A, Fonseca V, Giandhari J, Doolabh D, Pillay S, San EJ, Msomi N, Mlisana K, von Gottberg A, Walaza S, Allam M, Ismail A, Mohale T, Glass AJ, Engelbrecht S, Van Zyl G, Preiser W, Petruccione F, Sigal A, Hardie D, Marais G, Hsiao N-Y, Korsman S, Davies M-A, Tyers L, Mudau I, York D, Maslo C, Goedhals D, Abrahams S, Laguda-Akingba O, Alisoltani-Dehkordi A, Godzik A, Wibmer CK, Sewell BT, Lourenço J, Alcantara LCJ, Kosakovsky Pond SL, Weaver S, Martin D, Lessells RJ, Bhiman JN, Williamson C, de Oliveira T. 2021. Detection of a SARS-CoV-2 variant of concern in South Africa. Nature 592:438–443.

20. Faria NR, Mellan TA, Whittaker C, Claro IM, Candido D da S, Mishra S, Crispim MAE, Sales FCS, Hawryluk I, McCrone JT, Hulswit RJG, Franco LAM, Ramundo MS, de Jesus JG, Andrade PS, Coletti TM, Ferreira GM, Silva CAM, Manuli ER, Pereira RHM, Peixoto PS, Kraemer MUG, Gaburo N, Camilo C da C, Hoeltgebaum H, Souza WM, Rocha EC, de Souza LM, de Pinho MC, Araujo LJT, Malta FSV, de Lima AB, Silva J do P, Zauli DAG, Ferreira AC de S, Schnekenberg RP, Laydon DJ, Walker PGT, Schlüter HM, Dos Santos ALP, Vidal MS, Del Caro VS, Filho RMF, Dos Santos HM, Aguiar RS, Proença-Modena JL, Nelson B, Hay JA, Monod M, Miscouridou X, Coupland H, Sonabend R, Vollmer M, Gandy A, Prete CA, Nascimento VH, Suchard MA, Bowden TA, Pond SLK, Wu C-H, Ratmann O, Ferguson NM, Dye C, Loman NJ, Lemey P, Rambaut A, Fraiji NA, Carvalho M do PSS, Pybus OG, Flaxman S, Bhatt S, Sabino EC. 2021. Genomics and epidemiology of the P.1 SARS-CoV-2 lineage in Manaus, Brazil. Science 372:815–821.

21. Sheikh A, McMenamin J, Taylor B, Robertson C, Public Health Scotland and the EAVE II Collaborators. 2021. SARS-CoV-2 Delta VOC in Scotland: demographics, risk of hospital admission, and vaccine effectiveness. Lancet 397:2461–2462.

22. Martin DP, Weaver S, Tegally H, San JE, Shank SD, Wilkinson E, Lucaci AG, Giandhari J, Naidoo S, Pillay Y, Singh L, Lessells RJ, Gupta RK, Wertheim JO, Nekturenko A, Murrell B, Harkins GW, Lemey P, MacLean OA, Robertson DL, de Oliveira T, Kosakovsky Pond SL. 2021. The emergence and ongoing convergent evolution of the SARS-CoV-2 N501Y lineages. Cell 184:5189–5200.e7.

23. Mlcochova P, Kemp SA, Dhar MS, Papa G, Meng B, Ferreira IATM, Datir R, Collier DA, Albecka A, Singh S, Pandey R, Brown J, Zhou J, Goonawardane N, Mishra S, Whittaker C, Mellan T, Marwal R, Datta M, Sengupta S, Ponnusamy K, Radhakrishnan VS, Abdullahi A, Charles O, Chattopadhyay P, Devi P, Caputo D, Peacock T, Wattal C, Goel N, Satwik A, Vaishya R, Agarwal M, Indian SARS-CoV-2 Genomics Consortium (INSACOG), Genotype to Phenotype Japan (G2P-Japan) Consortium, CITIID-NIHR BioResource COVID-19 Collaboration, Mavousian A, Lee JH, Bassi J, Silacci-Fegni C, Saliba C, Pinto D, Irie T, Yoshida I, Hamilton WL, Sato K, Bhatt S, Flaxman S, James LC, Corti D, Piccoli L, Barclay WS, Rakshit P, Agrawal A, Gupta RK. 2021. SARS-CoV-2 B.1.617.2 Delta variant replication and immune evasion. Nature 599:114–119.

24. Cherian S, Potdar V, Jadhav S, Yadav P, Gupta N, Das M, Rakshit P, Singh S, Abraham P, Panda S, Team N. 2021. SARS-CoV-2 Spike Mutations, L452R, T478K, E484Q and P681R, in the Second Wave of COVID-19 in Maharashtra, India. Microorganisms 9:1542.

25. Delorey TM, Ziegler CGK, Heimberg G, Normand R, Yang Y, Segerstolpe Å, Abbondanza D, Fleming SJ, Subramanian A, Montoro DT, Jagadeesh KA, Dey KK, Sen P, Slyper M, Pita-Juárez YH, Phillips D, Biermann J, Bloom-Ackermann Z, Barkas N, Ganna A, Gomez J, Melms JC, Katsyv I, Normandin E, Naderi P, Popov YV, Raju SS, Niezen S, Tsai LT-Y, Siddle KJ, Sud M, Tran VM, Vellarikkal SK, Wang Y, Amir-Zilberstein L, Atri DS, Beechem J, Brook OR, Chen J, Divakar P, Dorceus P, Engreitz JM, Essene A, Fitzgerald DM, Fropf R, Gazal S, Gould J, Grzyb J, Harvey T, Hecht J, Hether T, Jané-Valbuena J, Leney-Greene M, Ma H, McCabe C, McLoughlin DE, Miller EM, Muus C, Niemi M, Padera R, Pan L, Pant D, Pe’er C, Pfiffner-Borges J, Pinto CJ, Plaisted J, Reeves J, Ross M, Rudy M, Rueckert EH, Siciliano M, Sturm A, Todres E, Waghray A, Warren S, Zhang S, Zollinger DR, Cosimi L, Gupta RM, Hacohen N, Hibshoosh H, Hide W, Price AL, Rajagopal J, Tata PR, Riedel S, Szabo G, Tickle TL, Ellinor PT, Hung D, Sabeti PC, Novak R, Rogers R, Ingber DE, Jiang ZG, Juric D, Babadi M, Farhi SL, Izar B, Stone JR, Vlachos IS, Solomon IH, Ashenberg O, Porter CBM, Li B, Shalek AK, Villani A-C, Rozenblatt-Rosen O, Regev A. 2021. COVID-19 tissue atlases reveal SARS-CoV-2 pathology and cellular targets. Nature 595:107–113.

26. Huang J, Hume AJ, Abo KM, Werder RB, Villacorta-Martin C, Alysandratos K-D, Beermann ML, Simone-Roach C, Lindstrom-Vautrin J, Olejnik J, Suder EL, Bullitt E, Hinds A, Sharma A, Bosmann M, Wang R, Hawkins F, Burks EJ, Saeed M, Wilson AA, Mühlberger E, Kotton DN. 2020. SARS-CoV-2 Infection of Pluripotent Stem Cell-Derived Human Lung Alveolar Type 2 Cells Elicits a Rapid Epithelial-Intrinsic Inflammatory Response. Cell Stem Cell 27:962–973.e7.

27. van der Vaart J, Lamers MM, Haagmans BL, Clevers H. 2021. Advancing lung organoids for COVID-19 research. Dis Model Mech 14:dmm049060.

28. Abo KM, Ma L, Matte T, Huang J, Alysandratos KD, Werder RB, Mithal A, Beermann ML, Lindstrom-Vautrin J, Mostoslavsky G, Ikonomou L, Kotton DN, Hawkins F, Wilson A, Villacorta-Martin C. 2020. Human iPSC-derived alveolar and airway epithelial cells can be cultured at air-liquid interface and express SARS-CoV-2 host factors. bioRxiv 2020.06.03.132639.

29. Krüger J, Groß R, Conzelmann C, Müller JA, Koepke L, Sparrer KMJ, Schütz D, Seufferlein T, Barth TFE, Stenger S, Heller S, Kleger A, Münch J. 2020. Remdesivir but not famotidine inhibits SARS-CoV-2 replication in human pluripotent stem cell-derived intestinal organoids. bioRxiv 2020.06.10.144816.

30. Nchioua R, Schundner A, Klute S, Noettger S, Zech F, Koepke L, Graf A, Krebs S, Blum H, Kmiec D, Frick M, Kirchhoff F, Sparrer KMJ. 2021. The Delta variant of SARS-CoV-2 maintains high sensitivity to interferons in human lung cells.

31. Li H, Liu T, Wang L, Wang M, Wang S. SARS-CoV-2 Delta variant infects ACE2low primary human bronchial epithelial cells more efficiently than other variants. Journal of Medical Virology n/a.

32. Thorne LG, Bouhaddou M, Reuschl A-K, Zuliani-Alvarez L, Polacco B, Pelin A, Batra J, Whelan MVX, Ummadi M, Rojc A, Turner J, Obernier K, Braberg H, Soucheray M, Richards A, Chen K-H, Harjai B, Memon D, Hosmillo M, Hiatt J, Jahun A, Goodfellow IG, Fabius JM, Shokat K, Jura N, Verba K, Noursadeghi M, Beltrao P, Swaney DL, Garcia-Sastre A, Jolly C, Towers GJ, Krogan NJ. 2021. Evolution of enhanced innate immune evasion by the SARS-CoV-2 B.1.1.7 UK variant.

33. Peacock TP, Sheppard CM, Brown JC, Goonawardane N, Zhou J, Whiteley M, Consortium PV, Silva TI de, Barclay WS. 2021. The SARS-CoV-2 variants associated with infections in India, B.1.617, show enhanced spike cleavage by furin.

34. Liu Y, Liu J, Johnson BA, Xia H, Ku Z, Schindewolf C, Widen SG, An Z, Weaver SC, Menachery VD, Xie X, Shi P-Y. 2021. Delta spike P681R mutation enhances SARS-CoV-2 fitness over Alpha variant. bioRxiv 2021.08.12.456173.

35. Kumar S, Thambiraja TS, Karuppanan K, Subramaniam G. 2021. Omicron and Delta Variant of SARS-CoV-2: A Comparative Computational Study of Spike protein.

36. Jacob A, Morley M, Hawkins F, McCauley KB, Jean JC, Heins H, Na C-L, Weaver TE, Vedaie M, Hurley K, Hinds A, Russo SJ, Kook S, Zacharias W, Ochs M, Traber K, Quinton LJ, Crane A, Davis BR, White FV, Wambach J, Whitsett JA, Cole FS, Morrisey EE, Guttentag SH, Beers MF, Kotton DN. 2017. Differentiation of Human Pluripotent Stem Cells into Functional Lung Alveolar Epithelial Cells. Cell Stem Cell 21:472–488.e10.

37. Jacob A, Vedaie M, Roberts DA, Thomas DC, Villacorta-Martin C, Alysandratos K-D, Hawkins F, Kotton DN. 2019. Derivation of self-renewing lung alveolar epithelial type II cells from human pluripotent stem cells. Nat Protoc 14:3303–3332.

38. Li H. 2013. Aligning sequence reads, clone sequences and assembly contigs with BWA-MEM. arXiv:13033997 [q-bio].

39. Li H, Handsaker B, Wysoker A, Fennell T, Ruan J, Homer N, Marth G, Abecasis G, Durbin R, 1000 Genome Project Data Processing Subgroup. 2009. The Sequence Alignment/Map format and SAMtools. Bioinformatics 25:2078–2079.

40. Grubaugh ND, Gangavarapu K, Quick J, Matteson NL, De Jesus JG, Main BJ, Tan AL, Paul LM, Brackney DE, Grewal S, Gurfield N, Van Rompay KKA, Isern S, Michael SF, Coffey LL, Loman NJ, Andersen KG. 2019. An amplicon-based sequencing framework for accurately measuring intrahost virus diversity using PrimalSeq and iVar. Genome Biology 20:8.

41. Nchioua R, Kmiec D, Müller JA, Conzelmann C, Groß R, Swanson CM, Neil SJD, Stenger S, Sauter D, Münch J, Sparrer KMJ, Kirchhoff F. 2020. SARS-CoV-2 Is Restricted by Zinc Finger Antiviral Protein despite Preadaptation to the Low-CpG Environment in Humans. mBio 11:16.

42. Groß R, Conzelmann C, Müller JA, Stenger S, Steinhart K, Kirchhoff F, Münch J. 2020. Detection of SARS-CoV-2 in human breastmilk. The Lancet https://doi.org/10.1016/S0140-6736(20)31181-8.

